# AFM-Fold: Rapid Reconstruction of Protein Conformations from AFM Images

**DOI:** 10.1101/2025.11.17.688836

**Authors:** Tsuyoshi Kawai, Yasuhiro Matsunaga

**Affiliations:** Graduate School of Science and Engineering, Saitama University; RIKEN Center for Computational Science

## Abstract

High-speed atomic force microscopy (HS-AFM) enables direct visualization of protein dynamics under near-physiological conditions, yet its intrinsic limitation to surface topography prevents atomic-level structural characterization. We present AFM-Fold, a generative AI-based framework that reconstructs three-dimensional protein conformations directly from AFM images. AFM-Fold combines a rotation-equivariant convolutional neural network, which extracts low-dimensional collective variables (CVs) from AFM images, with a guided diffusion process that generates conformations consistent with the inferred CVs. Using pseudo-AFM images of adenylate kinase, AFM-Fold accurately reproduced not only the open and closed conformations, but also intermediate states. Application to 159 experimental HS-AFM frames of the flagellar protein FlhA_C_ further demonstrated that AFM-Fold outperforms rigid-body fitting and captures time-correlated domain motions that reflect underlying conformational dynamics. AFM-Fold enables rapid, physically plausible structure estimation from individual AFM images, typically within one minute per frame, without relying on molecular dynamics simulations. This unified and computationally efficient pipeline opens a route to high-throughput structural analysis of HS-AFM movies.

## 1 Introduction

High-speed atomic force microscopy (HS-AFM) enables direct visualization of biomolecular dynamics in solution under near-physiological conditions, providing unprecedented insights into the relationship between biomolecular conformational dynamics and biological function [1, 2]. However, AFM measurements are inherently limited to surface topography, and the achievable spatial resolution is fundamentally constrained by the finite size of the cantilever tip [3, 4]. To address these limitations, there is a compelling need for computational modeling approaches that can integrate experimental AFM data with atomic-resolution three-dimensional structural models.

One approach is rigid-body fitting, in which template structures are selected from structural databases or molecular dynamics (MD) simulations and then exhaustively rotated and translated to find the best pose matching the AFM image [5, 6, 7, 8, 9, 10, 11, 12, 13]. Although powerful, this approach cannot account for conformational flexibility or changes; consequently, rigid-body fitting often fails to provide accurate structural models when the template library lacks conformations that match the AFM image. In real biological systems, the function of biomolecules is often coupled with conformational flexibility or dynamics. Thus, methods accounting for such conformational flexibility are required for analyzing and interpreting experimental AFM data.

In recent years, methods have been proposed that utilize MD or Monte Carlo (MC) simulation to generate conformations consistent with AFM images. For example, in flexible fitting [14, 15], the agreement with an AFM image is defined as an additional potential energy function *V*_AFM_, and MD simulations are driven to minimize it. In flexible fitting, sampling is often trapped in local minima, making optimization slow when separated by high barriers. If the global minimum is desired, prohibitively long simulations must be performed to ensure adequate sampling of the conformational space. To reduce the computational cost of MD, NMFF-AFM [16, 17] and AFMFit [18, 19] use a subset of normal modes to find conformations consistent with AFM images. In this case, however, the conformational space is fundamentally limited to capturing only harmonic fluctuations around a single reference structure. Another approach is to use a 3D Gaussian mixture model combined with MC sampling to estimate 3D low-resolution protein structures with conformational flexibility [20, 21]. In the field of AI-based structure modeling or prediction, sequence-to-structure models such as AlphaFold 2 [22, 23, 24], AlphaFold 3, and Boltz-1 [25] have demonstrated unprecedented precision in predicting the 3D fold of a protein. These models can be regarded as generative AI models that generate atomistic conformational ensembles from a given sequence. In principle, these generative AI models can generate not only specific peak structures present in the training data, but also diverse structural conformations. Moreover, it is known that the generation process can be navigated by applying conditioning or external forces during the generation stage. In the field of image generation, research has been conducted on navigating the generation process to bring generated images closer to specific targets. Recently, guiding the generative process of diffusion-based structure prediction models, including AlphaFold 3 and Boltz-1, has emerged as an approach aimed at generating targeted conformational ensembles. For example, in the context of NMR and X-ray crystallography data analysis [26], it has been demonstrated that imposing guided restraints during the generative process enables the generation of ensembles consistent with experimental observations. Similarly, for cryo-EM data [27], the incorporation of global or local density-map fitting as restraints has been proposed as a means of navigating the predicted conformations toward agreement with the experimental data.

In this study, we introduce AFM-Fold, which extends these ideas to AFM data analysis by employing structural restraints extracted from AFM images to guide the sampling trajectories of pre-trained protein structure generative models. Specifically, in the first step, a group-invariant convolutional neural network (CNN) [28] estimates low-dimensional collective variables (CVs, such as inter-domain distances) from AFM images, and then these estimated values are used as restraints to guide the generative process, enabling the rapid sampling of 3D conformations consistent with AFM images. We validated AFM-Fold in twin experiments. First, using adenylate kinase (AK, PDB IDs: 1AKE [29] and 4AKE [30]), we estimated the underlying CV values and accurately reconstructed conformations close to the ground truth. Second, we applied AFM-Fold to real experimental HS-AFM data from the flagellar protein FlhA_C_ (PDBID: 3A5I [31]) and confirmed that it predicts conformations more consistent with AFM images than the crystal structure fitted by rigid-body fitting calculations. Furthermore, we confirmed that AFM-Fold captures the time-correlated domain motions that reflect underlying conformational dynamics.

A distinctive feature of our framework, AFM-Fold, is that all steps—from training data generation, model training, and inference, to structure generation—are integrated within a single unified pipeline, without relying on time-consuming processes such as molecular dynamics simulations. This enables rapid reconstruction of structures from AFM images while reducing computational cost, allowing the observation of structural dynamics from hundreds of consecutive HS-AFM images. AFM-Fold provides a powerful framework for observing conformational transitions and functional mechanisms underlying HS-AFM images.

## 2 Related Work

Generative AI-based approaches employing diffusion models or flow models enable us to efficiently sample molecular conformations in a multiscale manner. For example, recent structure-generation foundation models, including AlphaFold 3 and Boltz-1, employ diffusion-based formulations. In particular, Boltz-2 [32] introduced *boltz-steering*, a framework that imposes a customized potential to navigate the sampling towards physically plausible conformations. These models largely follow a “one sequence–one structure” paradigm, which does not focus on producing diverse conformational ensembles or capturing conformational flexibility.

Several approaches have been proposed to address this problem. For example, the MSA subsampling technique [33] and its extensions [34, 35, 36] increase the diversity of multiple sequence alignments (MSAs) by masking parts of the MSA. Another powerful direction is the development of ensemble generative or MD surrogate models. An early contribution in this area is the Boltzmann generator [37], which was designed to learn and reproduce the Boltzmann distribution of small, specific peptides. More recently, models have been developed to generate diverse structural ensembles from sequences by learning from extensive MD simulation data across a wide range of proteins [38, 39, 40, 41, 42, 43, 44]. In these methods, obtaining sufficiently long MD trajectories for training requires substantial computational resources, and such models inevitably inherit the force-field biases present in the training data.

To mitigate this issue, recent studies have explored conditioning ensemble generative models on experimental measurements or physical prior knowledge. Examples include methods that generate NMR-consistent structural ensembles [26, 45, 46], and cryo-EM-consistent modeling using Boltz-1 and Chroma [27, 47]. Following this direction, AFM-Fold enables AFM-conditioned conformation generation, guiding AlphaFold 3’s prior distribution toward modeling more dynamic conformational states captured in HS-AFM images.

## 3 Methods

Our AFM-Fold framework leverages AlphaFold 3 (here, we used the open-sourced Protenix model [48], a PyTorch reimplementation of AlphaFold 3) to estimate the underlying 3D conformation from a given AFM image. To generate 3D structures consistent with AFM images using AlphaFold 3, we follow two main steps: (i) first, we estimate low-dimensional collective variables (CVs) for the target molecule from the given AFM image using a pre-trained CNN. For CVs, we use inter-domain distances, assuming multi-domain proteins as a typical case. (ii) Next, we use the estimated CVs to add restraints to AlphaFold 3 during structure generation, producing 3D structures that are consistent with the AFM image (see Fig. 1a).

**Figure 1.**
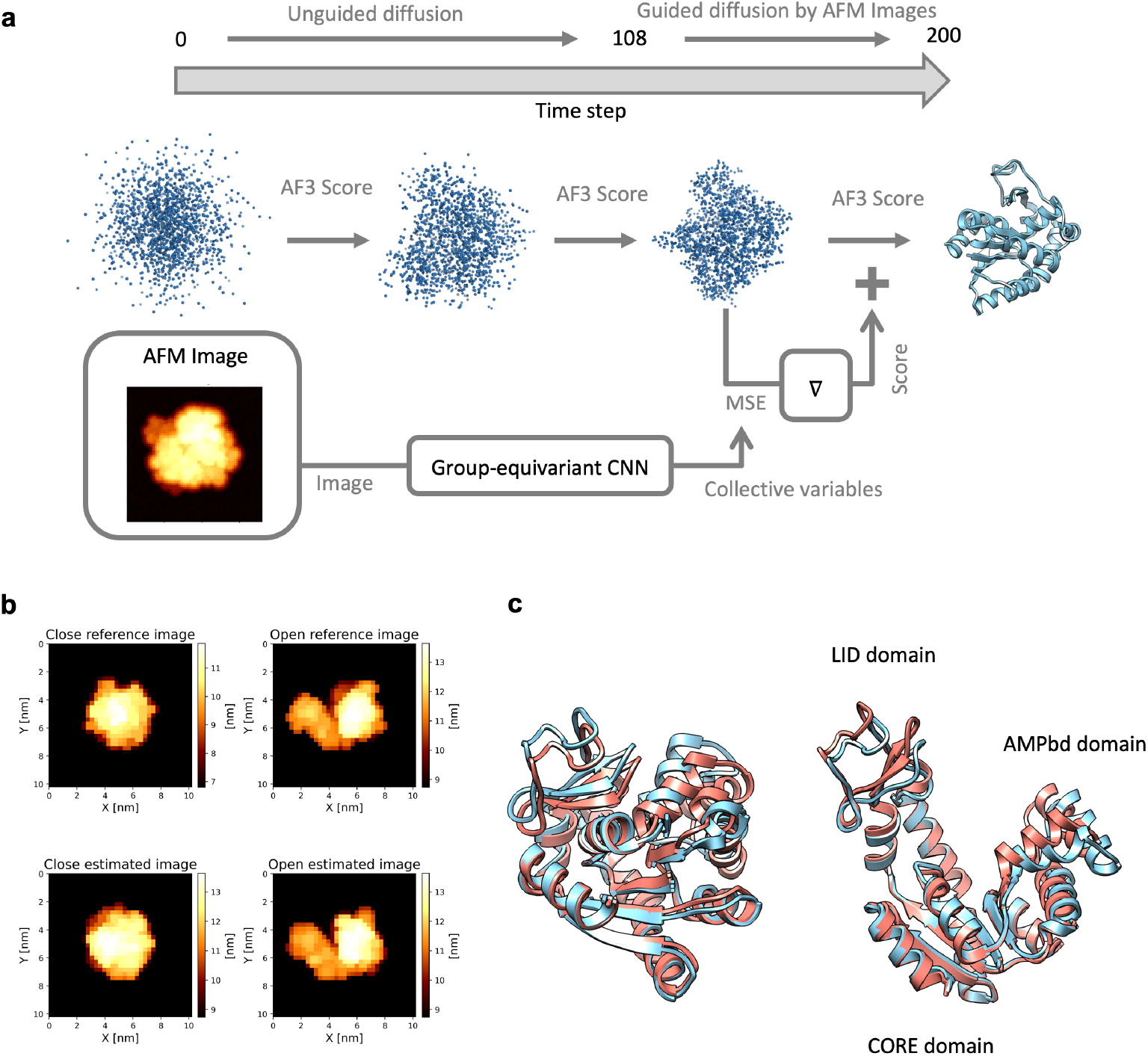
Schematic overview of AFM-Fold and outcome. AFM-based guidance navigates AlphaFold 3 and reproduces the closed and open states of AK. **(a)** Schematic diagram of AFM-Fold. The AFM image, shown at the lower left, serves as the input and is processed as follows: (1) coordinates in the inter-domain distance space are predicted from the AFM image by a group-invariant CNN (2) during the generative process, where the diffusion step *i* progresses discretely from 0 to 200, the mean squared error (MSE) between the predicted coordinates and the generated conformations is computed when *i >* 108; (3) the gradient of the MSE is used as a correction score to guide the generative process of AlphaFold 3. **(b)(c)** Results of structure prediction from pseudo-AFM images of AK. In (b), the upper panels show the reference pseudo-AFM images generated from the conformations in the PDB database (close: 1AKE, open: 4AKE). The lower panels show the reproduced pseudo-AFM images generated from the structures estimated by AFM-Fold. In (c), the blue structures (left: closed, right: open) represent the reference conformations, which were used to generate upper panels in (b). The red structures are the corresponding structures estimated by AFM-Fold.

In this section, we present our methodology in three parts. In Section 3.1, we summarize the group-equivariant CNN used to estimate the CVs from AFM images. In Section 3.2, we describe how we add restraints in the diffusion process of AlphaFold 3 using the estimated CVs. In Section 3.3, the overall computational protocol of our AFM-Fold framework is given. Finally, in **??**, we provide an overview of related work.

### 3.1 Group-equivariant CNNs

In AFM 2D image analysis, similar to single particle analysis of cryo-EM micrographs, the pose of the 3D structure is unknown. The alignment calculation between a 2D image and the 3D structural pose requires an exhaustive search and is computationally expensive. Moreover, if misalignment occurs, it leads to extreme deterioration in estimation accuracy. In the case of cryo-EM, the influence of misalignment could be mitigated on average by a vast number of micrographs, but in the case of HS-AFM, since we want to perform structural estimation for each single image to infer underlying conformational dynamics, misalignment cannot be tolerated. Therefore, we use a rotation-equivariant CNN model (*group-equivariant* CNN; g-CNN) [49, 50, 28, 51] that directly estimates structure-related CVs from 2D AFM images without performing alignment calculations.

In navigating the structure generation process of AlphaFold 3 to generate 3D structures consistent with AFM images, it would be possible to optimize by calculating image similarity at each step, as done in rigid-body fitting. In this case, higher-resolution image reproduction is expected, but on the other hand, we empirically found that it tends to produce structures trapped in local minima. This behavior arises because the image similarity landscape is highly multimodal and extremely sensitive to translational and in-plane rotational differences. Under a feasible amount of alignment search, inaccurate gradients are continuously applied, resulting in misleading forces during optimization. Therefore, instead of relying on image-to-image correlation, it was necessary to construct a comparison measure that is robust to in-plane rotations and translations of AFM images. In particular, to handle a large number of AFM frames, the similarity between the generated intermediate structures and the images needed to be computed efficiently.

With this motivation in mind, we summarize the theory of g-CNN [49, 50, 28, 51], and describe how it enables efficient estimation of CVs from AFM images. A standard CNN is naturally *translation equivariant* but not rotation equivariant. In contrast, g-CNN modifies feature indexing and kernel weight sharing so that the entire network becomes equivariant with respect to a chosen rotation group. Utilizing this architecture, the same physical state yields an identical representation regardless of its placement on the image plane, thereby reducing the need for extensive data augmentation and shortening training time. Consequently, we employ the g-CNN to estimate CVs from a single AFM image.

We model an AFM image as a single-channel (scalar) field *I*: ℝ^2^ *→* ℝ with pixel coordinate *x* = (*x*_1_, *x*_2_) *∈* ℝ^2^. In a standard CNN, intermediate feature maps are represented as functions *f*: ℝ^2^ *→* ℝ^*c*^. The input image is first mapped to a *c*-channel field by convolution with channel-specific filters 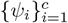:

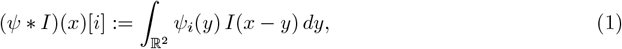

where * is convolution and *dy* denotes the area element. In practice, the integral is implemented as a finite sum over pixels; we keep the continuous notation for clarity.

Here, we consider the planar motion group 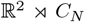, where *C*_*N*_ = *e, r*, …, *r*^*N−*1^ SO(2) is a finite rotation subgroup. For scalar or standard *c*-channel fields (without group indexing that we later explain), the action *π*(*t, g*) of translation *t ∈* ℝ^2^ and rotation *g ∈ C*_*N*_ for the field *u* is (*π*(*t, g*)*u*)(*x*):= *u* (*g*^*−*1^ (*x – t*)).

Because convolution commutes with translations, a standard CNN is translation-equivariant. For a single convolution with a (matrix-valued) kernel 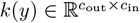,

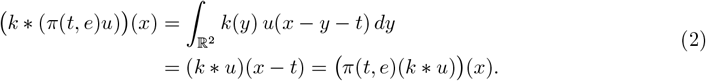

Since a typical activation *σ* is pointwise, *σ* is also translation-equivariant, *σ*(*π*(*t, e*)*u*) = *π*(*t, e*)*σ*(*u*) and the property propagates through the network. By contrast, the standard convolution is generally *not* rotation-equivariant.

The key idea of g-CNNs is to *lift* an image to a feature field indexed by group elements and to impose weight sharing consistent with the group action. Given a base filter *ψ*: ℝ^2^ *→* ℝ, define its rotated copies *ψ*_*h*_(*y*):= *ψ*(*h*^*−*1^*y*) for *h ∈ C*_*N*_, and set

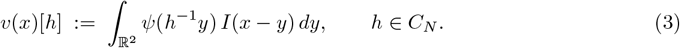

Thus *v*: ℝ^2^ *× C*_*N*_ *→* ℝ is a group-indexed (orientation-channel) field.

For such lifted fields, if the kernel satisfies the following constraint about sharing weights, *k*(*gy*)[*h, h*^*t*^] = *k*(*y*) [*g*^*−*1^*h, g*^*−*1^*h*^*t*^] for all *g, h, h*^*t*^ *∈ C*_*N*_, then the group convolution commutes with *π*(0, *g*):

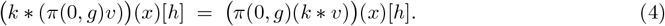

The proof can be found in Appendix D.1 of [28]. Hence, adding to the translation equivariance Eq. (2), we can construct CNNs equivariant to the chosen discrete rotation group *C*_*N*_. Channel-wise activations commute with the orientation relabeling, so equivariance to rotations (and translations) is preserved layer by layer.

A practical architecture uses multiple lifted blocks *v*_1_, …, *v*_*b*_; the overall action is the block-diagonal (direct-sum) representation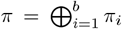. In AFM-Fold, because we require a vector invariant to rotations and translations, we apply group pooling and spatial pooling per block at last; i.e.,

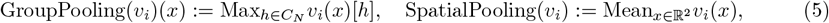

which yields features invariant to translations and rotations.

### 3.2 Navigating AlphaFold 3 with inter-domain distance restraints

Next, we describe a method for imposing a restraint on the generative process of AlphaFold 3 so that the CVs approach user-specified target values. Specifically, while AlphaFold 3 employs an Elucidated Diffusion Model (EDM) formulation [52], we compute the likelihood of the generated structure with respect to the specified CVs and add it to the original AlphaFold 3 score as a driving force that increases this likelihood. With this modified score, the generated structures become explicitly conditioned on the CVs.

Let *a* denote an amino acid sequence and *X* = (*x*_1_, …, *x*_*m*_) the 3D coordinates of all atoms, where *x*_*i*_ *∈* ℝ^3^. We write *X*_*ti*_ for the state at diffusion time *t*_*i*_ (*T > t*_1_ *> t*_2_ *>* …*> t*_*N*_ *>* 0). Sampling the generative process of AlphaFold 3 can be written as the stochastic EDM sampler:

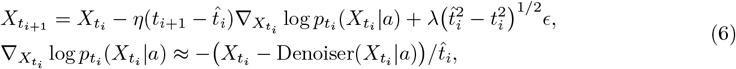

Here, 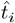denotes a time larger than *t*_*i*_, defined as 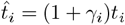, with a parameter *γ*_*i*_ that is set to 0.8 when *t*_*i*+1_ *>* 1.0, and 0 otherwise. The coefficients *η* and *λ* are hyperparameters called the *noise scale* and *step scale*, respectively, and are set to *η* = 1.5 and *λ* = 1.003. Denoiser(*·*|*a*) represents a denoiser conditioned by coevolutionary information about *a*.

Let *φ*_target_ *∈* ℝ^*D*^ denote the estimated coordinate in the CV space, where *D* is the dimension of the CVs. By Bayes’ rule, the conditional score decomposes as

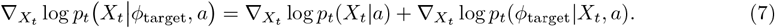

Therefore, we can bias the stochastic sampler by adding the guidance term:

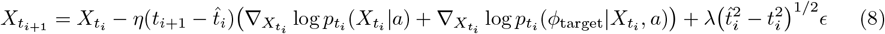

so that the process is guided toward *φ*(*X*_*tN*_) *≈ φ*_target_, where *φ*: ℝ^3*m*^ *→* ℝ^*D*^ is a differentiable mapping from structures to this feature space.

In this work, we replace the guidance score by the gradient of a squared-error loss:

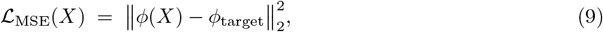

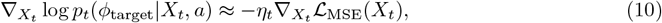

which is consistent with an isotropic Gaussian model *p*_*t (*_*φ*_object_|*X*_*t*_, *a))∝* exp 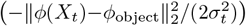, in which case 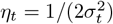. The optimal selection of *η*_*t*_ is analyzed in SI section S1.

### 3.3 AFM-Fold computational protocol

Our AFM-Fold framework consists of two main steps as described in the previous subsections: (i) estimation of CVs from AFM images using a g-CNN, and (ii) navigating the generative process of AlphaFold 3 using the estimated CVs. However, additional steps are still required for preparing training data for the g-CNN, training, inference, and evaluation. In this subsection, we describe these steps.

#### 3.3.1 Preparation of training data

To train the g-CNN to learn the relationship between AFM images and underlying 3D structures, we construct training data by generating pseudo-AFM images from 3D conformations and pairing them with ground-truth CV labels. In this study, these conformations are generated by applying diverse CV-based restraints to AlphaFold 3. This strategy offers two key advantages: (i) first, compared to other computational tools to generate 3D structures (such as MD simulations), the required computational resources are substantially reduced; (ii) second, this approach allows for sufficient sampling of intermediate conformations so that the g-CNN can learn to interpolate the structures in the conformational space. In order to detect smooth conformational transitions, the g-CNN requires training data not only for two boundary conformations (e.g., start and end conformations) but also for intermediate conformations continuously linking them. In the absence of such intermediates, the g-CNN is unable to accurately model continuous structural variability. By specifying the CV values continuously, we can generate diverse conformations that cover the conformational space.

##### Generating candidate conformations to broadly cover the CV space

We sampled diverse conformations by AlphaFold 3, broadly covering the CV space specifying the CV values with grids. To do this, first, we obtain a reference structure *X*_ref_ with AlphaFold 3 with no restraints. Then, target CV vectors 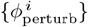 are placed on a grid around *φ*(*X*_ref_) such that, for each dimension *i*,

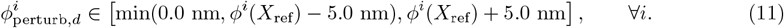

The grid used a step of 0.5 nm for each axis. Then, the generation with AlphaFold 3 is conducted using each 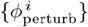 as the restraints for the navigating process.

##### Geometric sanitization

The candidate conformations generated in the previous step may contain geometric violations (e.g., steric clashes, backbone/side-chain outliers). Such conformations could bias the g-CNN by introducing unphysical patterns without valid conformational counterparts. To exclude such data from the training set, we applied thresholds to perform geometric sanitization, based on MolProbity scores (Table 1), excluding clear violations. This procedure constrains the g-CNN to perform inference within a conformational space devoid of geometric violations. The resulting set that passes this sanitization serves as a training set for training the g-CNN and for synthesizing pseudo-AFM images used in training.

**Table 1.**
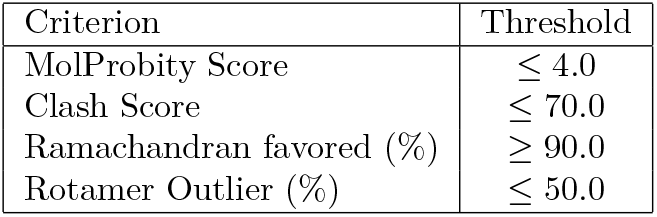
MolProbity score thresholds used for geometric sanitization.

| Criterion | Threshold |
| --- | --- |
| MolProbity Score | $\leq 4.0$ |
| Clash Score | $\leq 70.0$ |
| Ramachandran favored (%) | $\geq 90.0$ |
| Rotamer Outlier (%) | $\leq 50.0$ |

##### Pseudo-AFM image rendering

In rendering the pseudo-AFM, we used the morphology functions implemented in our previous work [4]. The specific parameter settings are summarized in Table 2. It should be noted that for the test case of AK, we used smaller values for the tip radius and finer resolution of pseudo-AFM images compared to typical experimental conditions. On the other hand, for FlhA_C_, since the precise tip geometry is unknown in experiments, the probe radius *r* was sampled uniformly within a range. Moreover, the height distribution of pseudo-AFM images was fitted to the real experimental images using `skimage.exposure.match_histograms`. For both AK and FlhA_C_, we used the inter-domain distances (distances between the centers of mass of C*α* atoms in the domains) as the CVs.

**Table 2.** Settings for training pseudo-AFM images.

|  | AK | FlhA<sub>C</sub> |
| --- | --- | --- |
| Lateral resolution [nm/pixel] | 0.3 | 0.98 |
| Pixel size [pixel × pixel] | $35 \times 35$ | $35 \times 35$ |
| Tip radius [nm] | $r \sim \text{Uniform}(0.3, 0.6)$ | $r \sim \text{Uniform}(2.0, 6.0)$ |
| Tip apex angle [degree] | $a \sim \text{Uniform}(10, 30)$ | $a \sim \text{Uniform}(10, 30)$ |
| Noise std. dev. [nm] | 0.0 | 0.5 |
| Histogram matching | — | ✓ |
| Dataset size [frames] | 5M | 5M |
| Collective variables | Inter-domain distances<br>(LID-CORE, CORE-AMPbd,<br>AMPbd-LID) | Inter-domain distances<br>(A<sub>C</sub>D<sub>1</sub>-A<sub>C</sub>D<sub>4</sub>, A<sub>C</sub>D<sub>2</sub>-A<sub>C</sub>D<sub>3</sub>,<br>A<sub>C</sub>D<sub>2</sub>-A<sub>C</sub>D<sub>4</sub>) |

#### 3.3.2 Training

Here, the training of the g-CNN can be regarded as supervised regression, where the task is to predict the CV values from a given AFM image. As the most straightforward loss function for supervised regression, we adopted the mean squared error (MSE) loss in the CV space:

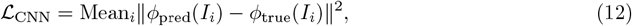

where *I*_*i*_ denotes the *i*-th input pseudo-AFM image, *φ*_pred_(*I*_*i*_) is the CV values predicted by the gCNN, and *φ*_true_(*I*_*i*_) is the corresponding ground-truth CV values. As the g-CNN architecture, we implemented a 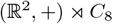-invariant CNN (see SI section S2 for the implementation details).

All models in this paper were trained on a single node equipped with an NVIDIA RTX A6000 GPU (48 GB memory). The training took approximately 2–3 days for both AK and FlhA_C_.

#### 3.3.3 Inference

The inference step consists of the estimation of CVs (inter-domain distances) from the AFM image by the g-CNN, and the navigation of the generative process of AlphaFold 3 using the estimated CVs. In this study, we require that the MSE of the g-CNN for the generated structures be below sub-Å, which is an important resolution for distinguishing different conformations for both AK and FlhA_C_.

The computation time for the inference step was extremely short compared to flexible fitting calculations using MD simulations. Indeed, inference time was less than one minute per image for both AK and FlhA_C_.

#### 3.3.4 Evaluation

To evaluate the accuracy of the estimated conformation, we computed the root mean square deviation (RMSD) between the estimated and the ground-truth conformations. Also, to evaluate how well the estimated conformation reproduces the given AFM image, we performed a rigid-body fitting, optimizing 3D rotation, translation, and tip radius. The optimization is evaluated through the correlation coefficient (c.c.) with the given AFM image, using Eq. (13) [14]. The pose achieving the maximum c.c. value across all rotations, translations and tip radii was taken as the optimal pose.

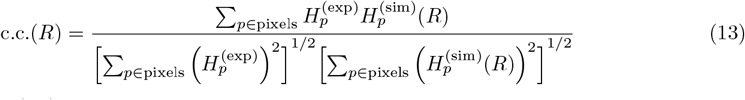

where 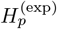and 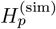 (*R*) denote the heights (i.e., the *z*-coordinates) of the given and generated image pixels (transformed by *R*), respectively, and the summation runs over all pixels *p*.

### 3.4 Molecular dynamics simulation

To verify the AFM-Fold framework, we performed MD simulations of AK and FlhA_C_. For AK, we performed an all-atom MD simulation to compare the distribution of conformational states sampled by MD with those generated by an AlphaFold 3 clone (Protenix [48]) for training the g-CNN. Simulations were conducted using the Amber ff14SB force field [53] with the TIP3P-FB water model and ions [54], starting from the open crystal structure (PDB ID: 4AKE). The simulations were carried out at 300 K and 1 atm using OpenMM [55], and the trajectory was extended to 450 ns to ensure sufficient sampling of the conformational space.

For FlhA_C_, due to computational time constraints, we did not conduct all-atom MD simulations. Instead, we generated conformational ensembles using a set of MD surrogate models: AlphaFlow [39] and BioEmu [44], as well as AlphaFold 2 with MSA subsampling [33] using ColabFold [24]. These generative approaches enabled us to obtain a diverse ensemble of conformations, facilitating a comprehensive comparison with the distribution generated by our AFM-Fold framework.

## 4 Results

### 4.1 Verification using pseudo-AFM images: adenylate kinase

adenylate kinase (AK) is a well-studied monomeric enzyme known for its large-scale conformational transitions. It is composed of three relatively rigid domains: the central CORE domain (residues 1–29, 68–117, and 161–214), the AMP-binding domain (AMPbd; residues 30–67), and the lid-like ATP-binding domain (LID; residues 118–167). Experimental and computational studies have suggested that upon ligand binding, the enzyme undergoes a transition from an inactive open conformation to an active closed conformation (see Fig. 1 (c)). Here, we evaluated how accurately AFM-Fold reconstructs 3D conformations of AK from artificially created pseudo “experimental” AFM images.

#### Quality of the conformational ensemble generated by AlphaFold 3

First, we checked the quality of the conformational ensemble generated by AlphaFold 3 used for the training of the g-CNN. Here, the conformations were generated by navigating AlphaFold 3 with CV values on a grid as described in Section 3.3.1. As the CVs, we used the inter-domain distances of all possible combinations of domain pairs (LID–CORE, CORE–AMPbd, and AMPbd–LID). To obtain physically plausible structures as accurately as possible via navigating to the target CVs, we systematically evaluated generated structures using various navigating schedules with different strengths and onset timings, and selected an optimal schedule strategy (see SI section S1 for details). Fig. 2 (b) shows the generated conformations projected to the LID–CORE, and CORE–AMPbd distances (colored by the AMPbd–LID distance), in comparison with Fig. 2 (a), which shows the projection of the conformations sampled by a 450 ns MD simulation. Compared with the MD simulation, the conformations generated by AlphaFold 3 adequately cover a broad conformational space of AK, including both closed (PDB ID: 1AKE) and open (PDB ID: 4AKE) crystal structures. This means that the g-CNN trained on this ensemble can learn the broad relationship between AFM images and the inter-domain distances of AK.

**Figure 2.**
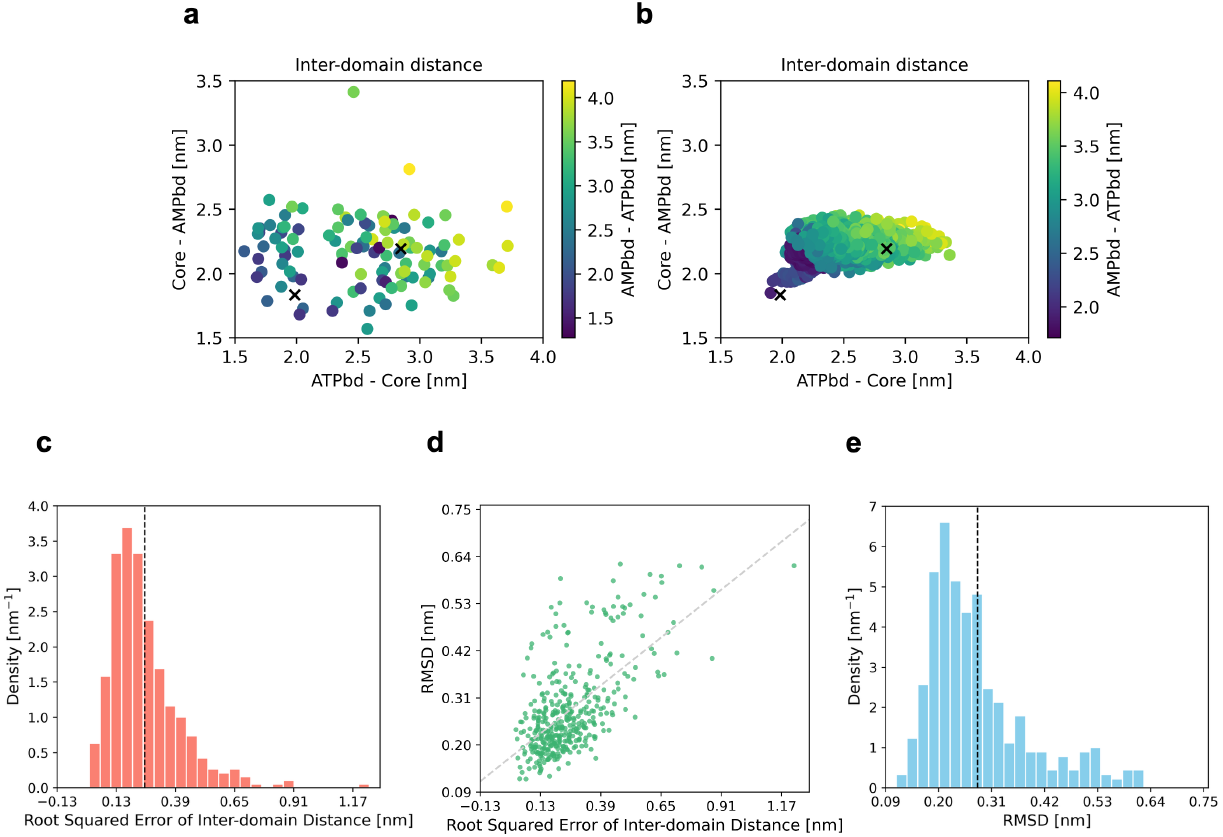
Training data and estimation accuracy of AFM-Fold. **(a)** Projection of training data into the inter-domain distance space. The two cross marks represent the open (4AKE) and closed (1AKE) crystal structures from PDB, respectively. **(b)** Projection of a 450 ns MD simulation trajectory into the inter-domain distance space. **(c)** Distribution of the root squared error in the inter-domain distance space between the g-CNN’s estimations and the ground-truth. The inter-domain distance was estimated by the trained g-CNN from pseudo-AFM images, which were generated from randomly selected molecular dynamics simulation conformations. The average of the root squared errors is indicated by the broken line (0.254 nm). **(d)** Scatter plot of the root squared error in the inter-domain distance space and the heavy-atom RMSDs from the ground-truth conformation. The heavy-atom RMSD was computed between the conformation generated with AlphaFold 3 navigating using the predicted CV values, and the ground-truth conformation. **(e)** Density of the heavy-atom RMSDs. The average of the RMSD is indicated by the broken line (0.281 nm).

#### Accuracy of collective variables estimated by the g-CNN

Next, we investigated whether the trained g-CNN accurately estimates CV values from given pseudo-AFM images. Here, we estimated CV values from pseudo-AFM images emulated using randomly chosen conformations from the MD simulation trajectory. The pseudo-AFM images were generated with the settings described in Table 2. Fig. 2 (c) shows the distribution of the root squared error in the CV space between the estimated values with the trained g-CNN and the ground-truth values. With the exceptions of some outlier errors greater than 0.500 nm, the trained g-CNN achieves the mean root squared error of 0.254 nm, which is enough resolution to capture intermediate conformational states of AK.

#### Accuracy of conformations estimated by AFM-Fold

Then, we investigated the accuracy of the estimated conformations of our AFM-Fold framework. As a demonstration, we first estimated the closed and open conformations of AK from the pseudo-AFM images emulated from the closed and open crystal structures (the images are shown in Fig. 1 (b)), respectively. Fig. 1 (c) shows the estimated structures by the protocol of AFM-Fold (i.e., g-CNN and AlphaFold 3 navigation). The RMSD between the ground-truth and reconstructed conformations was 0.216 nm for the closed form and 0.176 nm for the open form, respectively. Considering that the RMSD between the closed and open crystal structures is 0.715 nm, AFM-Fold estimation is accurate enough to identify the two states. The c.c. between the AFM images created from the estimated conformations and the given “experimental” pseudo-AFM image is 0.997 for the closed form and 0.998 for the open form, respectively (here, Eq. (13) was used to compute the c.c.).

We then evaluated the RMSDs of the conformations estimated from the pseudo-AFM images emulated from randomly chosen conformations from the MD simulation trajectory. Fig. 2 (e) shows the distribution of the RMSDs between the ground-truth and estimated conformations. Overall, the RMSD values are low and, aside from a small number of outliers, most RMSDs are distributed around the mean value of approximately 0.281 nm. Fig. 2 (d) shows the scatter plot of the RMSD and the root squared errors of the predicted CVs between the ground-truth and estimated conformations. Although a reasonable correlation is observed between the two measures, some scatter was observed in RMSD at similar error levels; that is, even with comparable CV errors, the RMSDs can vary. Upon close inspection of the generated structures, we found that this arises because a given set of current CVs (inter-domain distances) does not always uniquely determine the domain arrangement. Specifically, either hinge-bending (open–close transition) or shear-like (twisted) domain motions can yield similar inter-domain distances, despite resulting in structurally distinct conformations. This degeneracy in the CV–structure mapping explains the occurrence of RMSD outliers even when the predicted CV values are accurate.

#### Correlation coefficient and robustness against noise

To investigate the applicability of our approach to real experimental data, we next focused on two aspects: (i) the development of evaluation metrics that can be computed without ground-truth structures and (ii) the assessment of robustness in structure estimation accuracy against noise. As for the first aspect, following the previous study [14], we propose to use the c.c. between the given experimental image and a pseudo-AFM image created by rigid-body fitting of the estimated structure to the experimental image, maximizing the c.c. values. Fig. 3 (a) shows the density of the c.c. between the “experimental” images (pseudo-AFM images created from randomly selected structures from the MD data) and pseudo-AFM images created by rigid-body fitting of the estimated structures. With the exceptions of some outlier values, the distribution is centered around the average value of 0.998, indicating that the estimated structures are in good agreement with the given images. Fig. 3 (b) shows the scatter plot of the c.c. and the RMSD. A clear negative correlation is observed between the c.c. and the RMSD, suggesting that, even when ground-truth structures are unavailable and RMSD cannot be computed, as is often the case with real experimental data, the comparison of c.c. values provides a useful indication of the reliability of the estimated structures.

**Figure 3.**
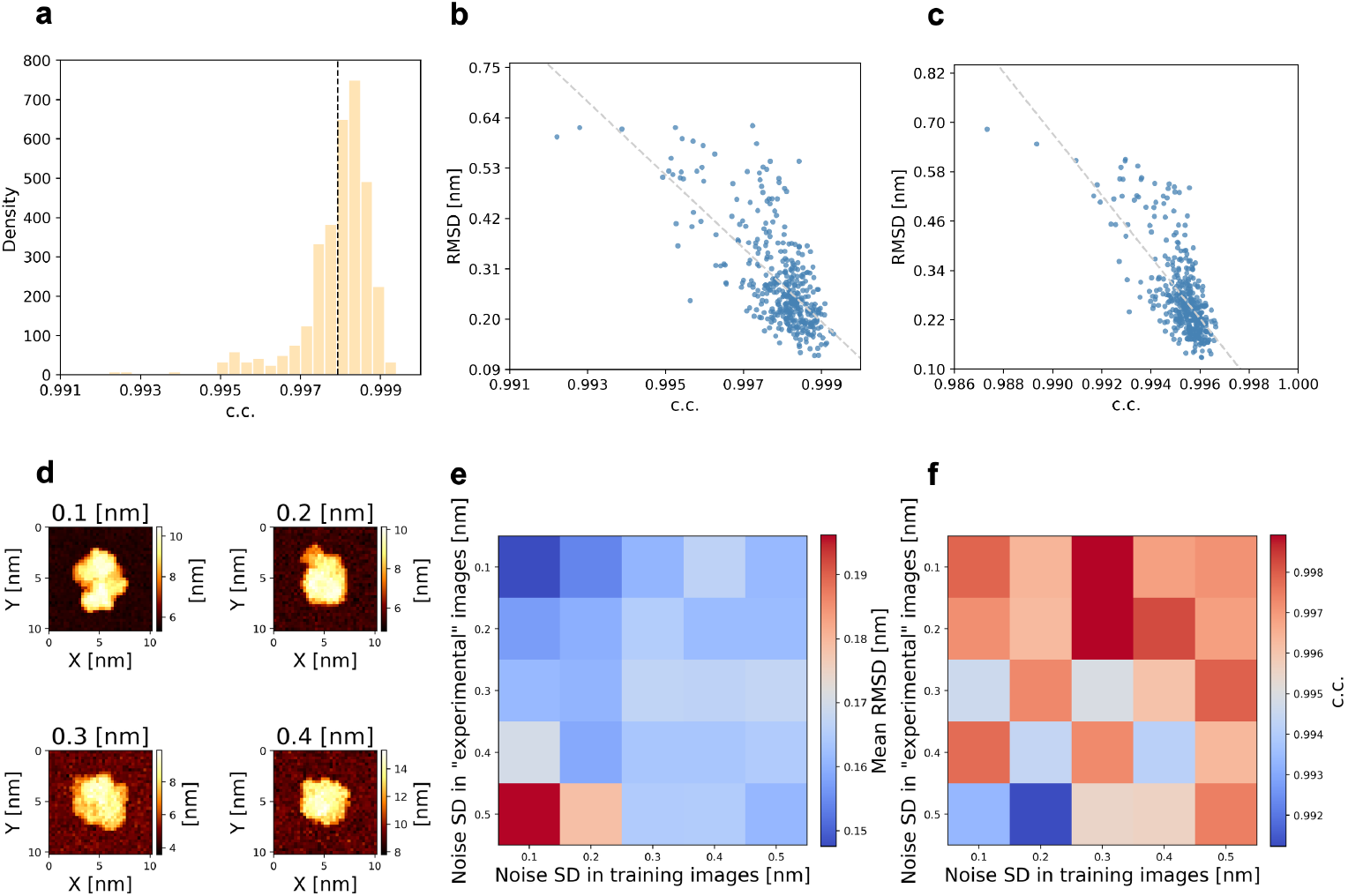
Correlation coefficients between the given “experimental” images and the pseudo-AFM images generated from the estimated structures, and robustness against noise. Panels **(a)** and **(b)** present the evaluations under noise-free conditions, whereas panels **(c–f)** evaluate the robustness of the estimations against noise. **(a)** Distribution of correlation coefficients (c.c.) between the given images (“experimental” pseudo-AFM images) and pseudo-AFM images, each of which was generated by a rigid-body fitting of an estimated conformation to the reference image, maximizing the c.c. value. The average of c.c. is indicated by the broken line (0.998). **(b)** Scatter plot of the c.c. and the RMSD. **(c)** Scatter plot of the c.c. and the RMSD using “experimental” pseudo-AFM images contaminated by Gaussian noise with a standard deviation of 0.3 nm. **(d)** Examples of “experimental” pseudo-AFM images used for estimating structures at different noise levels in the investigation of noise robustness. The images were generated from randomly selected structures from the molecular dynamics simulation trajectory and then adding Gaussian noise with various standard deviations. **(e)** Heat map of the mean RMSD between the estimated structures and the ground-truth structures for various noise levels in the training and “experimental” pseudo-AFM data. Each mean RMSD was computed for 20 independent structure estimations. **(f)** Heat map of the mean c.c. between the rigid-body fitted pseudo-AFM images and the “experimental” pseudo-AFM images under the same noise levels as in panel (e).

Next, we focused on the second aspect, the assessment of robustness in structure estimation accuracy against noise. So far, we have used noise-free pseudo-AFM images for both the training and inference stages. To assess the robustness of the estimation against noise, we first added noise to “experimental” pseudo-AFM data in the inference stage using Gaussian noise with a standard deviation of 0.3 nm (which is a typical noise level for HS-AFM images [14]). Fig. 3 (c) shows the scatter plot of the c.c. and the RMSD using the noisy “experimental” pseudo-AFM images. Still, a clear negative correlation is observed between the c.c. and the RMSD, suggesting that the c.c. values would work even when noisy real experimental data are used.

Furthermore, for systematic evaluation of the robustness against various noise levels, we systematically varied the noise levels added to both the training and the “experimental” pseudo-AFM data. Fig. 3 (d) illustrates examples of pseudo-AFM images at different noise levels, and Fig. 3 (e) shows a heat map of the mean RMSD between the estimated and ground-truth structures over the range of noise levels. Notably, we found that adding noise to the “experimental” pseudo-AFM images consistently led to substantial reductions in prediction accuracy. In contrast, adding noise to the training data alone had a relatively minor effect on the estimation accuracy; indeed, the best performance was achieved when noise-free training data were used. This indicates that, while robustness to noise in the inputs (“experimental” images) is limited, the presence or absence of noise in the training data does not substantially affect accuracy, with noise-free training data providing the most reliable results. Fig. 3 (f) shows a heat map of the mean c.c. between the rigid-body fitted pseudo-AFM images and the “experimental” pseudo-AFM images under the same noise levels as in Fig. 3 (e).

### Application to real experimental AFM images: a flagellar protein FlhA_C_

FlhA is a key component of the flagellar protein export system, and its C-terminal domain, FlhA_C_, helps transport proteins by assisting their binding to the exporter. Structurally, it comprises four domains (A_C_D_1_, A_C_D_2_, A_C_D_3_, and A_C_D_4_) and a linker to the transmembrane domain (FlhA_TM_) (Fig. 4 (a)). Previous studies using X-ray crystallography and mutational analysis indicate that hinge motions among these domains influence transport activity and motility [31].

**Figure 4.**
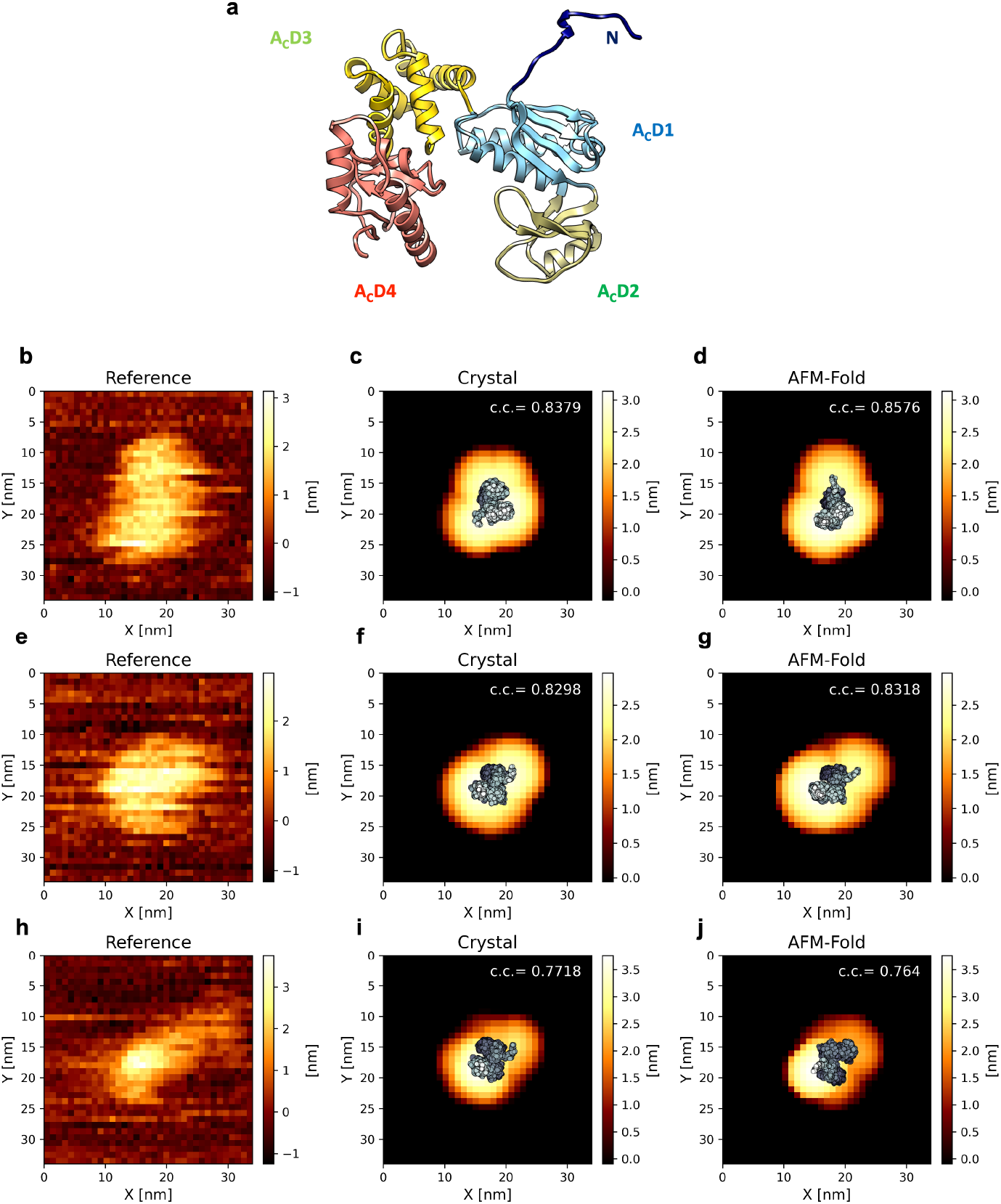
FlhA_C_ structure and estimated structures compared with the crystal structure. Crystal structure of FlhA_C_ (PDB ID: 3A5I). The four domains are colored as follows: dark blue, linker; sky blue, A_C_D_1_; green, A_C_D_2_; yellow, A_C_D_3_; and orange, A_C_D_4_. **(b,e,h)** Experimental AFM images selected from real experimental data. (b) shows the case where *c*.*c*._AFMFold_ *− c*.*c*._Crystal_ is maximal, panel (e) shows the case where it is at the median value, and panel (h) shows the case where it is minimal. **(c,f,i)** Crystal structures fitted to the corresponding experimental AFM images through rigid-body fitting maximizing the c.c. value. Pseudo-AFM images generated from them are shown in the background. **(d,g,j)** Estimated structures by AFM-Fold, fitted to the corresponding experimental AFM images through rigid-body fitting maximizing the c.c. value. Pseudo-AFM images generated from them are shown in the background.

Here, to evaluate the applicability of AFM-Fold to real AFM data [56], we estimated the conformations of FlhA_C_ using AFM-Fold from 159 HS-AFM images of the FlhA_C_ monomer. It should be noted that, whereas conventional flexible fitting methods for capturing structural changes typically require hours to days of computation per AFM image even with a coarse-grained model [14], AFM-Fold allows estimation of conformations from each AFM image in approximately one minute on a single GPU (considering training time, total computation time is less than the conventional flexible fitting methods). This enabled, for the first time, the processing of as many as 159 AFM images in this study.

#### Training of the g-CNN

Conformations for training the g-CNN were generated by navigating AlphaFold 3 with CV values on a grid as described in Section 3.3.1. As the CVs, we used the interdomain distances of domain pairs, A_C_D_1_–A_C_D_4_, A_C_D_2_–A_C_D_3_, and A_C_D_2_–A_C_D_4_. The generated ensembles are shown with their MolProbity scores in Fig. S5. The g-CNN was trained on the training data to learn the relationship between AFM images and the CVs.

#### Evaluating estimated conformations by AFM-Fold

Fig. 4 shows the estimated structures of three AFM images selected from the total 159 AFM images. We performed rigid-body fitting of the estimated structures to the real AFM image by exhaustively rotating and translating and finding the pose that maximizes the c.c. value, and then their poses and pseudo-AFM images were drawn for comparison with the real AFM images. As a comparison, we performed the same fitting procedure for the crystal structure (PDB ID: 3A5I) of FlhA_C_ to the three AFM images (also shown in Fig. 4). Although it is difficult to distinguish by visual inspection alone, the three rows respectively represent cases where AFM-Fold performed most clearly better than rigid fitting using the crystal structure (top), a typical, median case (middle), and a case where AFM-Fold did not outperform the crystal-structure-based rigid fitting (bottom). In most cases, the c.c. value between the pseudo-AFM image and the real AFM image was consistently higher for the estimated structure than for the crystal structure, although this difference is often subtle. This is because the overall c.c. value is strongly influenced by whether the molecule is correctly centered in the image, whereas the contribution of finer structural features—such as subtle surface height variations—is comparatively minor, as shown previously for AK. Cases in which AFM-Fold failed to reproduce the reference images were often associated with poor image resolution or measurement noise, which made the apparent size of the molecule in the real AFM image very small.

Next, we evaluated the estimated conformations by AFM-Fold using the total 159 images. Fig. 5 (a) shows the comparison of the c.c. values between the estimated structures by AFM-Fold and the crystal structure with the real AFM images. Again, in almost all cases, the c.c. value for the estimated structure was higher than that for the crystal structure. Interestingly, some of the c.c. values of AFM-Fold take higher values when the estimated structures have more open conformations in Fig. 5 (a) (measured by the distance between A_C_D_2_ and A_C_D_4_, which serves as a useful indicator to distinguish between open and closed conformations). This might suggest that the flexibility of the inter-domain motions of FlhA_C_ is well captured by AFM-Fold.

**Figure 5.**
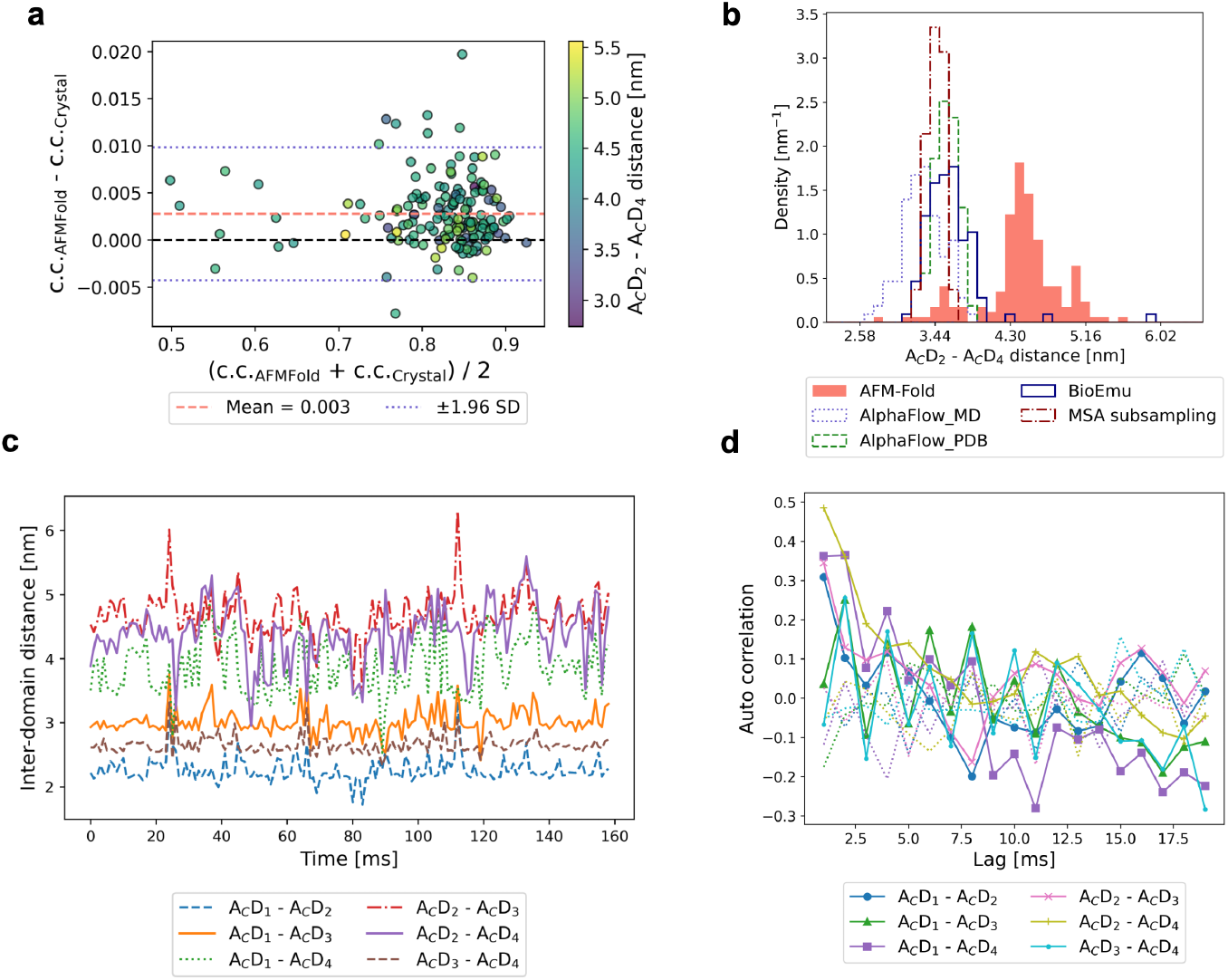
Analysis of the estimated structures by AFM-Fold. **(a)** Comparison of correlation coefficients between pseudo-AFM images generated from the AFM-Fold estimations and the experimental AFM images (159 frames) with those of the crystal structure. The *x*-axis represents the mean of c.c._AFM Fold_ and c.c._Crystal_, and the *y*-axis shows their difference (c.c._AFM_ *−* _Fold_*−* c.c._Crystal_). (b)Distribution of the inter-domain distances of the AFM-Fold estimations compared with AI-based structure generation models. **(c)** Time series of inter-domain distances estimated by AFM-Fold for the consecutive 159 frames of the experimental AFM images. **(d)** Auto-correlation functions of the inter-domain distance as a function of time lag. The solid line shows the results from AFM-Fold estimations, while the dashed line corresponds to temporally shuffled data.

To quantify the flexibility of the inter-domain motions captured by AFM-Fold, we computed the distribution density of the inter-domain distance between A_C_D_2_ and A_C_D_4_ for all the estimated structures (Fig. 5(b)). For comparison, we also generated conformations using AI-based structure generation models, AlphaFlow [39], BioEmu [44], and AlphaFold 2 with MSA subsampling [33]. These methods are expected to approximate long-time behaviors of FlhA_C_ that are not accessible by brute-force MD simulations. Interestingly, compared with these models, AFM-Fold tends to estimate more open conformational states. There are three possible explanations for why AFM-Fold tends to estimate more open conformational states than the AI-based structure generation models. First, HS-AFM captures molecular dynamics on a timescale of 1 ms per frame, which may reflect longer-timescale motions than those considered by AlphaFlow or BioEmu, thereby enabling the observation of more open structures. Second, interactions between the molecule and the stage during AFM imaging may stabilize conformations with larger surface areas in contact with the stage, thus favoring more open structures. Third, errors in the structural estimation by AFM-Fold could result in an overrepresentation of highly open conformations with large entropy. These factors together may account for the observed tendency toward open states in our estimations.

To verify the third possibility (large estimation errors in AFM-Fold), we investigated the time-series of the estimated structures by AFM-Fold (Fig. 5 (c)). Since the 159 AFM images are a set of consecutive frames as time-series data, we can expect some temporal correlations in the estimated structures if the estimation errors are small. Among the inter-domain distances shown in Fig. 5 (c), the distance between A_C_D_2_ and A_C_D_4_ or A_C_D_1_ and A_C_D_4_ serves as a key indicator distinguishing between open and closed conformations. Examination of the time-series reveals oscillatory behavior, with the distance fluctuating between larger (open) and smaller (closed) values over time, Importantly, this temporal pattern suggests the presence of time correlation in the estimated structures. Despite the fact that AFM-Fold estimates each frame independently without considering temporal information, the observed time correlation in estimated inter-domain distances implies that the estimation errors are smaller than the scale of the observed fluctuations.

The temporal correlation of the inter-domain distances was quantified by the auto-correlation function (Fig. 5 (d)). The solid lines show the auto-correlations from AFM-Fold estimations, while the dashed lines show the auto-correlations from temporally shuffled surrogate data (i.e., the frame order of the inter-domain distances time-series is shuffled). In particular, the auto-correlation for the temporally shuffled data remains low (within ±0.2) across all lag times, indicating little to no time correlation as expected for randomly ordered data. In contrast, the auto-correlation for the AFM-Fold estimations shows a pronounced positive correlation for lag times up to 2.0 ms and exhibits larger negative values for lag times beyond 10 ms, resulting in oscillatory behavior. Overall, these results demonstrate that AFM-Fold is effective for capturing the molecular conformational dynamics from the time-series of HS-AFM images.

## 5 Discussion

In this work, we have proposed a computational protocol, AFM-Fold, to predict the 3D conformation of biomolecules from AFM images. AFM-Fold enables rapid and accurate conformation estimation compared to previous approaches. AFM-Fold extracts features from AFM images with a group-invariant (symmetry-aware) CNN, reducing computational cost in two ways. First, it reduces the amount of required training data, because symmetry-aware representations learn efficiently from limited images [28]. Second, it shortens inference time: guiding AlphaFold 3 requires computing *p*_*t*_(*φ*_object_ |*X*_*t*_, *a*); replacing this with *L*_MSE_(*X*_*t*_) keeps the computational cost per structure low. In the estimations of this study, per-frame estimation finished within one minute on a single GPU. It has the potential to serve as a powerful tool for interpreting HS-AFM movies comprised of hundreds of frames in both structural and dynamic aspects.

A limitation of AFM-Fold is that it requires a priori definition of CVs as features to be estimated from the AFM image. If it is challenging to define appropriate CVs a priori, one could first perform structural sampling using methods such as AlphaFlow [39], BioEmu [44], or AlphaFold 2 with MSA subsampling [33], and then identify meaningful CVs by applying dimensionality reduction techniques like PCA to extract important features. The AFM-Fold framework is general, and as long as the chosen CVs are differentiable with respect to atomic Cartesian coordinates and can be used for guiding AlphaFold 3, a wide variety of CV types can be incorporated. For example, although we focused on inter-domain distances as CVs in this work, other CVs, such as residue-level distance restraints, can be used to enable AFM-guided reconstruction of finer-grained motions without a significant increase in computational cost.

Another limitation of AFM-Fold is that it infers conformations from a single AFM image and currently does not quantify predictive uncertainty due to image noise or molecular orientation. Extending the framework to estimate uncertainty would make the predictions easier to interpret [57]. Moreover, AFM image sequences constitute time-series data in which poses change only slightly between adjacent frames; incorporating temporal dependencies—for example, with TimeSformer [58] or non-local networks [59]—could further improve fidelity.

Finally, in this work we navigated AlphaFold 3’s sampling only using AFM images; consequently, neither structural stability nor explicit physical priors are incorporated. In the future, conditioning the prior (such as Boltz-2’s custom potentials [32]) of an ensemble generative model should enable integrated modeling that unifies experimental measurements, physical priors, and computational inference.

## 6 Data availability

The code of AFM-Fold is publicly available at http://github.com/matsunagalab/afmfold. Notebooks, scripts, and datasets to reproduce the results of this paper are publicly available at Zenodo https://zenodo.org/records/17714204 [60].

## 7 Acknowledgements

We appreciate Tohru Minamino (The University of Osaka) and Noriyuki Kodera (Kanazawa University) for kindly providing the experimental AFM image data used in this work. This work was supported by JSPS KAKENHI (Grant number: 23H03412), and partly supported by MEXT as “Program for Promoting Researches on the Supercomputer Fugaku” (Development and application of large-scale simulation-based inferences for biomolecules JPMXP1020230119) and used computational resources of supercomputer Fugaku provided by the RIKEN Center for Computational Science (Project IDs: hp230209, hp240215, hp250233).

