## Supporting Information for "AFM-Fold: Rapid Reconstruction of Protein Conformations from AFM Images"

**Supporting Materials**

### S1 Navigating scheduling

We investigated how the navigating schedule $\eta_t$ in Eq. (8) affects the navigating of AlphaFold 3 structure generation. The conditional generation process we used in this study is formulated as:

$$X_{t_{i+1}} = X_{t_i} - \eta(t_{i+1} - \hat{t}_i)\big(\nabla_{X_{t_i}} \log p_{t_i}(X_{t_i}|a) - \eta_{t_i} \nabla_{X_{t_i}} \mathcal{L}_{\mathrm{MSE}}(X_{t_i})\big) + \lambda\sqrt{\hat{t}_i^2 - t_i^2}\,\epsilon, \tag{S1}$$

where $i$ propagates from 1 to $N$ and $\eta_t$ dominates the strength of the navigation.

Prior work commonly used simple schedules——e.g., a constant value [1] or a mid-trajectory sigmoid [2]—— but, to our knowledge, there has not been a systematic quantification of how these choices affect generated structures. Here, across different navigating schedules, we evaluated the trade-off between the accuracy of following the target CV values and the physical plausibility of the generated structures, and found an optimal navigating parameter set.

**Navigating family and design of the sweep.** We defined navigating schedules $\eta_t$ from the sigmoidal family defined in Eq. (S2). Specifically, we vary two hyperparameters that determine *when* navigating turns on ($\tau_{\mathrm{start}}$) and *how strong* it becomes at the end ($y_{\max}$). For the two hyperparameters, we select five candidate values each: $\tau_{\mathrm{start}}$ is chosen at evenly spaced points in $[0.4, 0.9]$, and $y_{\max}$ is chosen at evenly spaced points in $[0.6, 1.6]$. By evaluating all combinations of these choices, we obtained $5 \times 5 = 25$ distinct schedule settings in total. For each setting we generated five independent conformations and computed the averages of evaluation metrics of them.

$$\eta_{t_i} = \frac{y_{\max}}{2}\frac{\|\delta\|_2}{\|g\|_2}\left(1 + \frac{\tanh(\alpha(\tau_i - \beta)/2)}{\tanh(\alpha\beta/2)}\right), \qquad \tau_i = \frac{i/N - \tau_{\mathrm{start}}}{1 - \tau_{\mathrm{start}}},$$
$$\delta = \nabla_{X_{t_i}} \log p_{t_i}(X_{t_i}), \quad g = -\nabla_{X_{t_i}} \mathcal{L}_{\mathrm{MSE}}(X_{t_i}), \quad \alpha = 10.0, \quad \beta = 0.5. \tag{S2}$$

**Systems and collective variables** Here, we describe the choice of protein system and the collective variables (CVs) on which restraints are applied. The way restraints are imposed can in principle depend on factors such as protein size or the specific CVs chosen, but our goal is to provide a concrete reference point that can serve as a guideline when users consider how to set the guidance strength.

In this experiments, we investigated the relationship between the guidance strength and the physical plausibility of the generated structures for both AK (214 residues) and $\mathrm{FlhA_C}$ (346 residues). The choice of CVs used for applying the guidance followed the same setup as in the main part—that is, LID–CORE, CORE–AMPbd, and AMPbd–LID for AK, and $\mathrm{A_CD_1}$ –$\mathrm{A_CD_4}$ and $\mathrm{A_CD_2}$ –$\mathrm{A_CD_3}$ for $\mathrm{FlhA_C}$. As a reference, under guidance-free conditions, AlphaFold 3 predicts structures with inter-domain distances of (2.85 nm, 2.19 nm, 3.45 nm) for AK, and (3.47 nm, 4.38, nm) for $\mathrm{FlhA_C}$. We conducted two types of experiments for these proteins: (i) **in-distribution targets**, which correspond to physically plausible inter-domain distances observed in MD simulations. Specifically, (2.41 nm, 2.01, nm, 2.68, nm) was chosen for AK and (2.5 nm, 4.5 nm) for $\mathrm{FlhA_C}$. (ii) **out-of-distribution targets**, which represent physically implausible inter-domain distances not observed in MD simulations. In this case, (1.21 nm, 1.51, nm, 0.53, nm) was chosen for AK and (2.0 nm, 3.0, nm) for $\mathrm{FlhA_C}$.

Across both the in-distribution and out-of-distribution targets, three clear patterns emerge.

- First, there is a dividing line in the MSE landscape, approximately connecting $(\tau_{\mathrm{start}}, y_{\max}) =$ (0.65, 0.60) and (0.73, 1.60) (see Fig. S2 (b, h) and Fig. S3 (b, h)). On the right side of this line (i.e., with later $\tau_{\mathrm{start}}$ and smaller $y_{\max}$), the MSE between the target CVs and the generated conformation's CVs rises sharply.

- Second, the MolProbity-score contours are roughly circular, centered near $(\tau_{\mathrm{start}}, y_{\max}) =$ (0.40, 1.60), rather than linear (see Fig. S2 (a, g) and Fig. S3 (a, g)).

- Third, combining these two observations, we find that starting the guidance early (small $\tau_{\mathrm{start}}$) and keeping the restraint weak (low $y_{\max}$) enables the application of constraints without substantially compromising the physical plausibility of the generated structures.

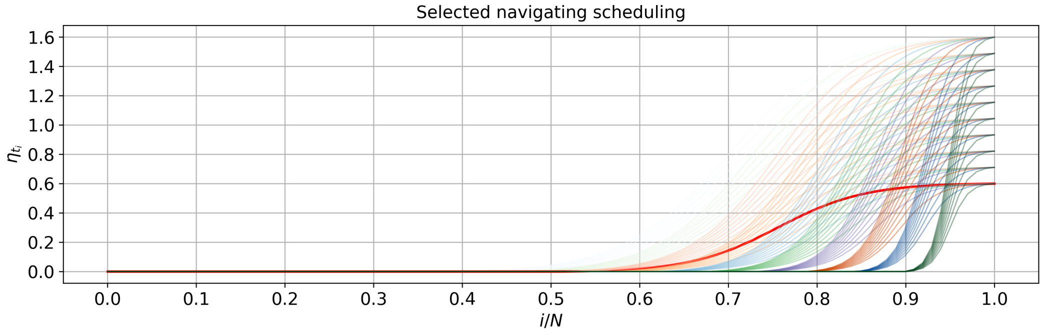


Figure S1: All navigating schedulings evaluated in this study. The setting used in our main experiments, $\tau_{\text{start}} = 0.6$ and $y_{\max} = 0.6$, is highlighted with a thick red line.

Overall, we observed the expected trade-off: stronger guidance improves agreement with the target, but tends to degrade structural plausibility. At the same time, we obtained an additional insight: rather than applying strong restraints only in the final stages of the reverse process, it is more effective to apply weak restraints from the early stages. In fact, regions with low MSE consistently appear in the area of $\hat{t}_{\text{start}} \lessapprox 0.6$ (see Fig. S2 (b,h) and Fig. S3 (b,h)), and among them, regions with low MolProbity scores are located where $\hat{y}_{\max}$ is relatively small (see Fig. S2 (a,g) and Fig. S3 (a,g)).

Based on these observations, in our study we adopt $\hat{t}_{\text{start}} = 0.6$ and $y_{\max} = 0.6$ that allow effective guidance without substantially degrading structural quality.

### S2 Model Implementation Details

**Algorithm S1** Forward function of $(\mathbb{R}^2, +) \rtimes C_8$ invariant CNN

**Input:** $I \in \mathbb{R}^{B \times D \cdot N \times H \times W}$, $D = 1$, $N = 8$, $H = W = 35$;
channels = [3,6,6,12,12,8], kernel_size = [7,5,5,5,5,5],
$N_{\text{hidden-layers}} = 3$, $D_{\text{hidden}} = 64$, $D_{\text{out}} = N_{\text{domain-pairs}}$
**Output:** $\mathbf{D} \in \mathbb{R}^{B \times D_{\text{out}}}$
$I \leftarrow$ R2Conv($I$; in $= D \cdot N$, out $=$ channels[0] $\cdot N$, $k =$ kernel_size[0]) $\rightarrow$ InnerBatchNorm $\rightarrow$ ReLU
**for** $i = 1 \rightarrow$ len(channels) $- 1$ **do**
  $I \leftarrow$ R2Conv($I$; in $=$ channels[$i-1$]$\cdot N$, out $=$ channels[$i$]$\cdot N$, $k =$ kernel_size[$i$])
  $I \leftarrow$ InnerBatchNorm($I$) $\rightarrow$ ReLU($I$)
  **if** $i \bmod 2 = 0$ **then**
    $I \leftarrow \text{PointwiseAvgPoolAntialiased}\left(I;\ \sigma = 0.66,\ \text{stride} = \begin{cases} 2 & \text{if size odd} \\ 1 & \text{if size even} \end{cases}\right)$
$I' \leftarrow$ GroupPooling($I$)    $I' \in \mathbb{R}^{B \times \text{channels}[-1] \times H' \times W'}$
$I'' \leftarrow$ PointwiseAvgPool($I'$)    $I'' \in \mathbb{R}^{B \times \text{channels}[-1] \times 1 \times 1}$
$\mathbf{z} \leftarrow$ Flatten($I''$)    $\mathbf{z} \in \mathbb{R}^{B \times \text{channels}[-1]}$
$\mathbf{h} \leftarrow$ Linear($\mathbf{z}$; in $=$ channels[$-1$], out $= D_{\text{hidden}}$) $\rightarrow$ BatchNorm1d $\rightarrow$ ELU    $\mathbf{h} \in \mathbb{R}^{B \times D_{\text{hidden}}}$
**for** $j = 1 \rightarrow N_{\text{hidden-layers}} - 1$ **do**
  $\mathbf{h} \leftarrow$ Linear($\mathbf{h}$; in $= D_{\text{hidden}}$, out $= D_{\text{hidden}}$) $\rightarrow$ ReLU($\mathbf{h}$)    $\mathbf{h} \in \mathbb{R}^{B \times D_{\text{hidden}}}$
$\mathbf{D} \leftarrow$ Linear($\mathbf{h}$; in $= D_{\text{hidden}}$, out $= D_{\text{out}}$)    $\mathbf{D} \in \mathbb{R}^{B \times D_{\text{out}}}$
**return D**

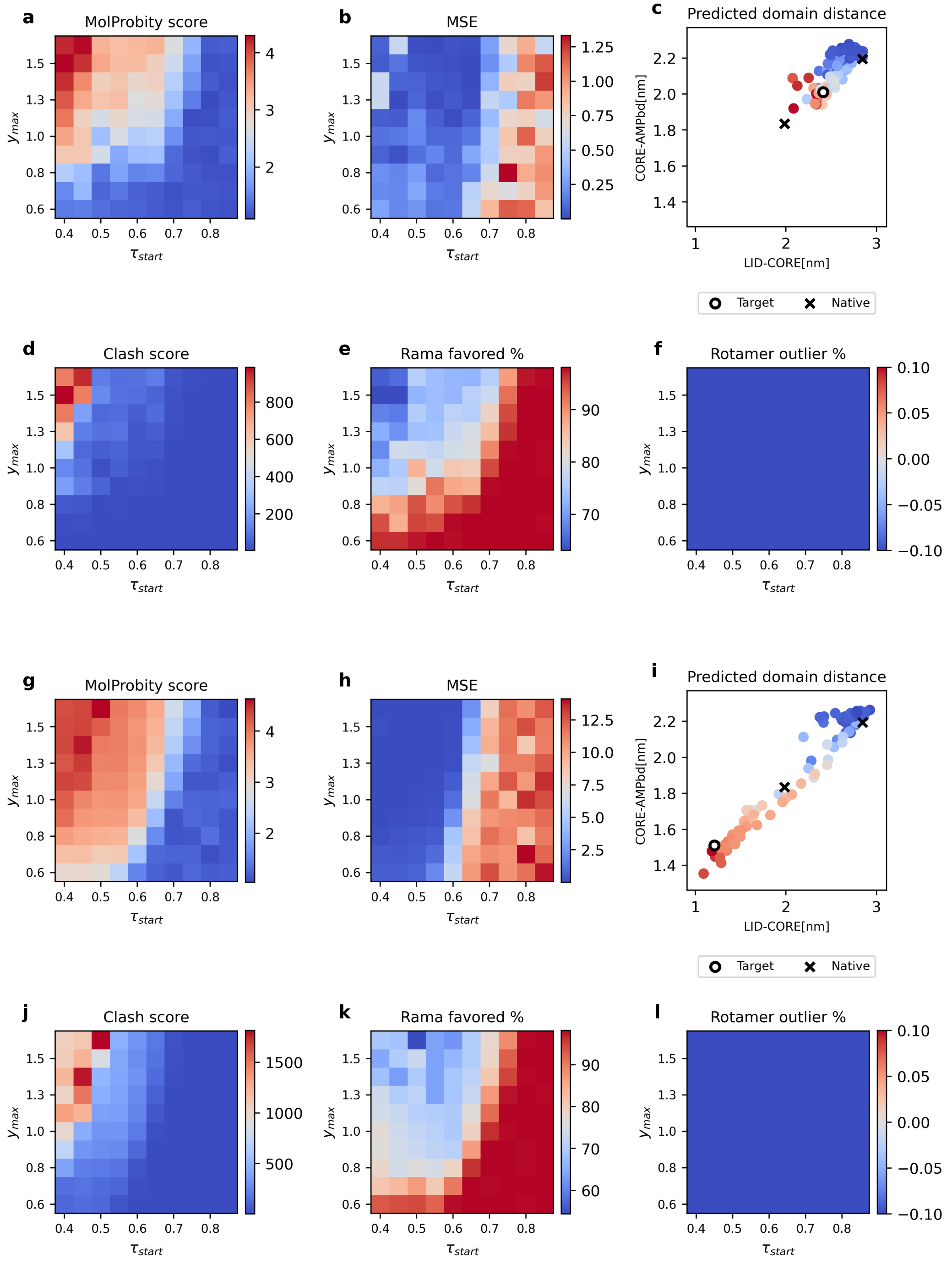


Figure S2: **Results for AK**. The upper 6 panels ((a)–(f)) shows results for the in-distribution target, and the lower 6 panels ((g)–(l)) shows results for the out-of-distribution target. **(a), (g)** Average MolProbity scores. **(b), (h)** Mean squared error (MSE). **(c), (i)** The inter-domain distance of estimated conformations (25 points, each one represents average inter-domain distance of corresponding schedule setting), with point colors representing the MolProbity scores. The cross point means that of the crystal structures (PDBID: 4AKE (open) and 1AKE (close)), and circle point means that of the target (every estimated structures were generated navigated towards this point). **(d), (e), (f), (j), (k), (l)** Evaluation metrics obtained during MolProbity scoring: clash score, Ramachandran favored rate (%), and rotamer outlier rate (%).

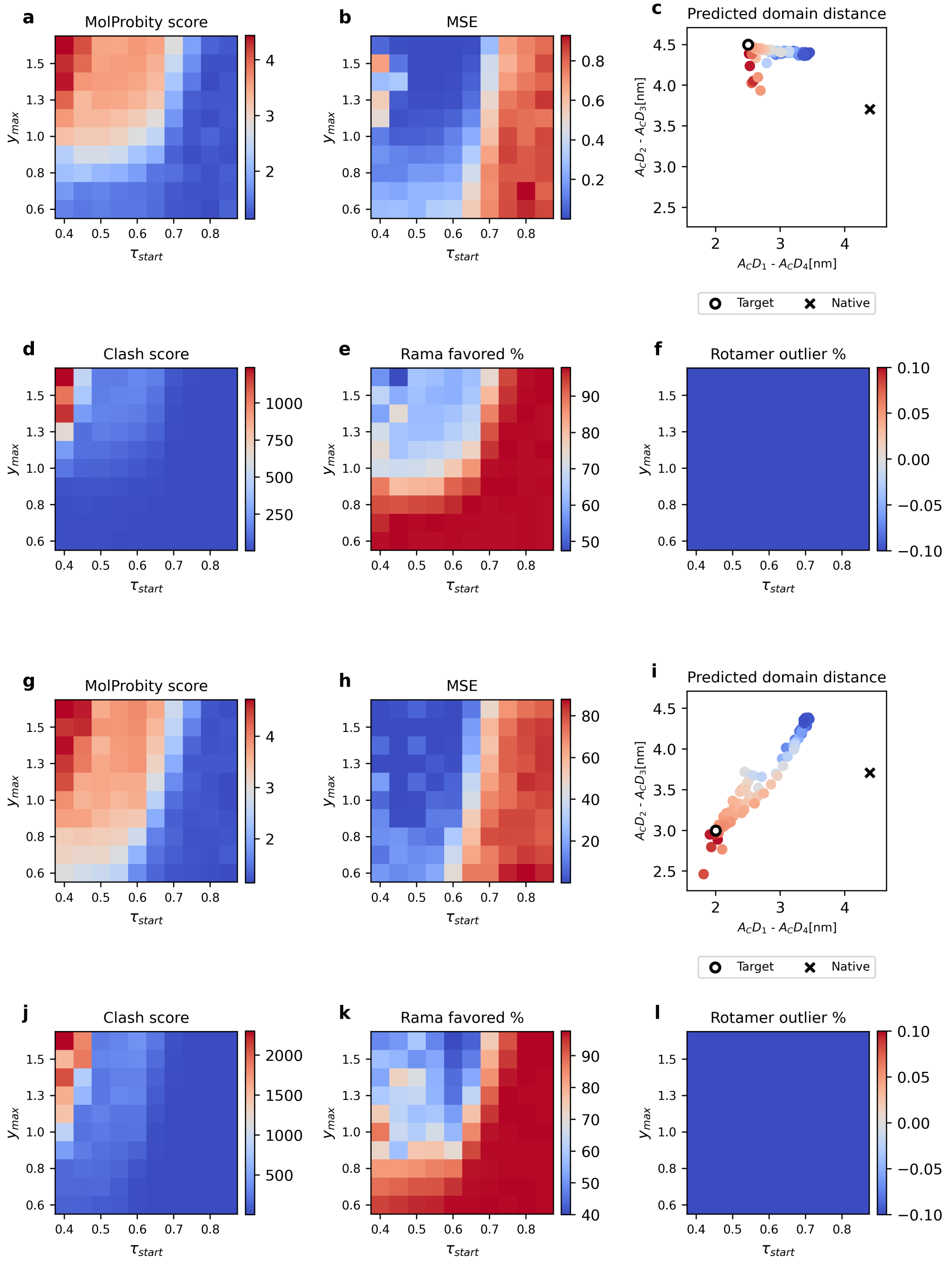


Figure S3: **Results for FlhA$_\mathbf{C}$.** As before, the upper 6 panels ((a)–(f)) shows results for the in-distribution target, and the lower 6 panels ((g)–(l)) shows results for the out-of-distribution target. **(a), (g)** Average MolProbity scores. **(b), (h)** Mean squared error (MSE). **(c), (i)** The inter-domain distance of estimated conformations, with point colors representing the MolProbity scores. The cross point means that of the crystal structure (PDBID: 3A5I), and circle point means that of the target. **(d), (e), (f), (j), (k), (l)** Evaluation metrics obtained during MolProbity scoring: clash score, Ramachandran favored rate (%), and rotamer outlier rate (%).

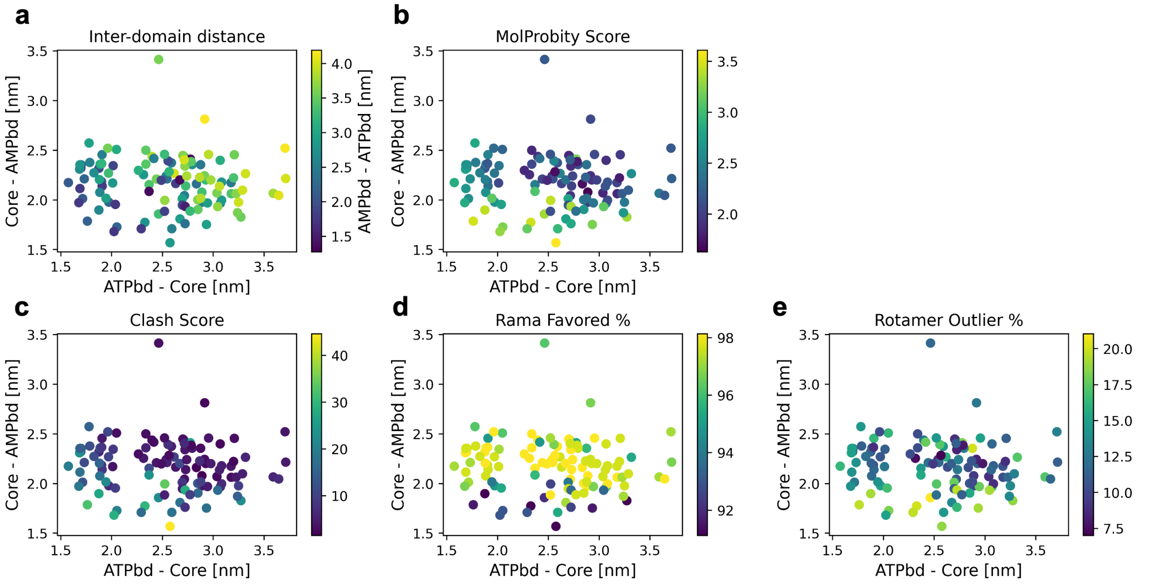


Figure S4: **Selected candidate conformations for AK and MolProbity evaluation for them**. **(a)** Projection of the candidate conformations into the inter-domain distance space. The following panels shows **(b)** MolProbity score, **(c)** Clash score, **(d)** Ramachandran favored rate (%), and **(e)** Rotamer outlier rate (%) of the candidate conformations.

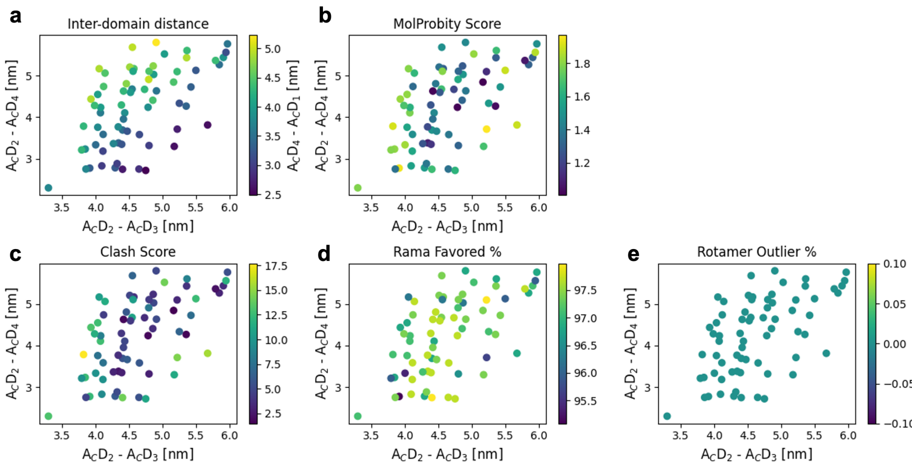


Figure S5: **Selected candidate conformations for FlhA$_C$ and MolProbity evaluation for them**. **(a)** Projection of the candidate conformations into the inter-domain distance space. The following panels shows **(b)** MolProbity score, **(c)** Clash score, **(d)** Ramachandran favored rate (%), and **(e)** Rotamer outlier rate (%) of the candidate conformations.
